# Profiling and structural analysis of cardenolides in two species of Digitalis using liquid chromatography coupled with high-resolution mass spectrometry

**DOI:** 10.1101/864959

**Authors:** Baradwaj Gopal Ravi, Mary Grace E. Guardian, Rebecca Dickman, Zhen Q. Wang

## Abstract

Plants of the *Digitalis* genus contain a cocktail of cardenolides commonly prescribed to treat heart failure. Cardenolides in *Digitalis* extracts have been conventionally quantified by high-performance liquid chromatography yet the lack of structural information compounded with possible co-eluents renders this method insufficient for analyzing cardenolides in plants. The goal of this work is to structurally characterize cardiac glycosides in fresh-leaf extracts using liquid chromatography coupled with tandem mass spectrometry (LC/MS/MS) that provides exact masses. Fragmentation of cardenolides is featured by sequential loss of sugar units while the steroid aglycon moieties undergo stepwise elimination of hydroxyl groups, which distinguishes different aglycones. The sequence of elution follows diginatigenin→digoxigenin→gitoxigenin→gitaloxigenin→digitoxigenin for cardenolides with the same sugar units but different aglycones using a reverse-phase column. A linear range of 0.8-500 ng g^−1^ has been achieved for digoxigenin, *β*-acetyldigoxin, and digitoxigenin with limits of detection ranging from 0.09 to 0.45 ng g^−1^. A total of 17 cardenolides have been detected with lanatoside A, C, and E as major cardenolides in *Digitalis lanata* while 7 have been found in *Digitalis purpurea* including purpurea glycoside A, B, and E. Surprisingly, glucodigifucoside in *D. lanata* and verodoxin and digitoxigenin fucoside in *D. purpurea* have also been found as major cardenolides. As the first MS/MS-based method developed for analyzing cardenolides in plant extracts, this method serves as a foundation for complete identification and accurate quantification of cardiac glycosides, a necessary step towards understanding the biosynthesis of cardenolide in plants.

## 1. Introduction

Digoxin extracted from the woolly foxglove (*Digitalis lanata*) has been frequently used to treat systolic heart failure and atrial fibrillation [1]. It is on the World Health Organization’s Model List of Essential Medicines recommended for all health systems, rendering it one of the most commonly prescribed plant-based pharmaceuticals [1,2]. Advances in organic synthesis have achieved total synthesis of digoxin, but cultivating the host plant remains the only cost-effective option for subsequent clinical use [3,4]. Digoxin belongs to a large family of cardiac glycosides found in the genus of *Digitalis* and other plant lineages as defense chemicals against insect herbivores [5]. The composition of cardenolides varies greatly among different species of foxglove and is also influenced by climate and geographic locations [6]. Structurally, cardenolides are steroid glycosides with a pentacyclic aglycon and zero to four monosaccharides linked by the *β*-O-glycosyl bond to the C3 of the steroid (Fig. 1). Fresh *Digitalis* leaves contain primary cardenolides with three digitoxose and one glucose moiety. The terminal glucose is readily hydrolyzed away during the drying process of fresh leaves, generating secondary cardenolides such as digoxin. The aglycones of *Digitalis* cardenolides are distinguished from other phytosteroids due to the unique C17*β* substitution with a γ-butyrolactone ring and a 14 *β*-hydroxylation. Different aglycons of *D. lanata* cardenolides (digitoxigenin, gitoxigenin, digoxigenin, diginatigenin, gitaloxigenin) vary in both the numbers and positions of hydroxyl groups. In gitaloxigenin, the 16-hydroxyl group is further modified to a formate ester.

**Fig. 1.**
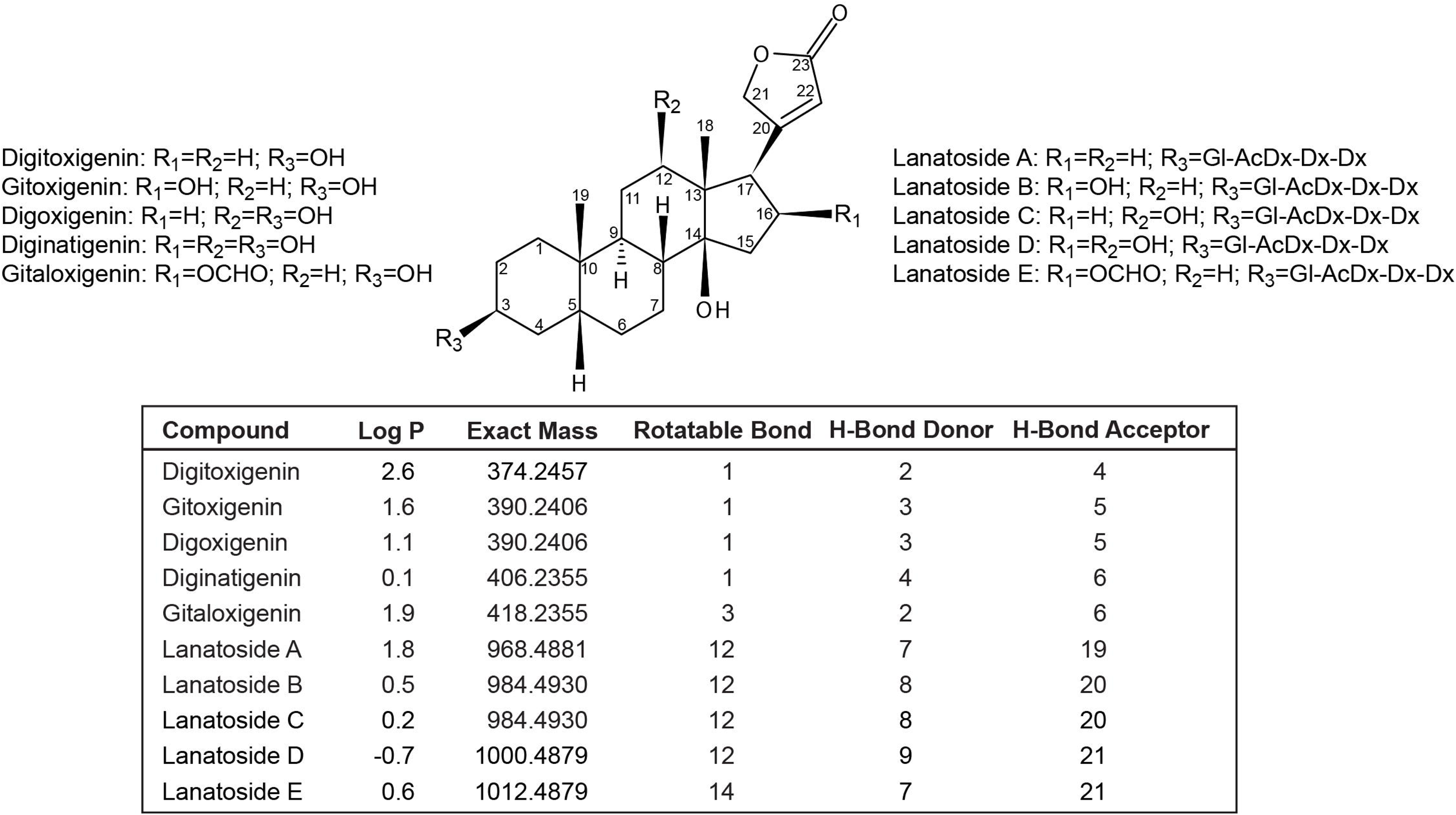
Chemical structures and physicochemical properties of primary cardenolides and their corresponding aglycones in *D. lanata*. Gl: glucose unit; Dx: digitoxose unit; AcDx: acetyldigitoxose unit. Log P: octanol-water partition coefficient. H-bond, hydrogen bond. Rotatable bonds are defined as any single non-ring bond, bound to a nonterminal heavey (non-hydrogen) atom.

Intriguing progress has been made in the taxonomy and *in vitro* propagation of *Digitalis* in the last two decades, but progress in structural elucidation and analytical methods for cardenolides *in planta* is lagging in development [6]. High-performance liquid chromatography (HPLC) coupled with diode-array detection remains the most used method for the identification and quantification of plant cardenolides. The HPLC method developed by Wiegrebe et. al. in 1993 remains the most comprehensive profiling of cardenolides in *D. lanata* [7]. However, HPLC-based methods are not ideal for cardenolides due to these substances’ low extinction coefficients in the UV range. Also, structural information cannot be obtained and co-eluents from the complex plant matrix interfere with quantification. Previous liquid chromatography tandem mass spectrometry (LC/MS/MS) developed for cardiac glycosides mainly analyzed blood or urine samples, and only digoxin and few of its metabolic derivatives were monitored [8-16]. Therefore, a highly sensitive and selective LC/MS method is necessary for structural validation and accurate quantification of cardenolides in crude plant extracts.

Here we report for the first time a high-resolution mass spectrometry (HRMS) method for profiling and structural characterization of cardiac glycosides from fresh leaves of *D. lanata* and *D. purpurea*. The HRMS method measures atomic mass units (amu) to four decimal places, allowing the determination of the molecular formula of unknown compounds within 5 ppm of their exact masses [17]. Subsequent collision-induced dissociation (CID) provides fragments containing structural information of the targeted cardenolides. We find that cardenolides sequentially lose sugar units through the breakage of glycosidic bonds and that different aglycones can be distinguished by patterns of stepwise dehydration during CID. 17 cardenolides in fresh leaves of *D. lanata* and 7 in *D. purpurea* have been identified with structural information for both the aglycones and the sugars. This HRMS method lays the foundation for the comprehensive characterization of cardiac glycosides in various plant species as a step towards understanding the biosynthesis of these medicinally important plant natural compounds.

## 2. Material and Methods

### 2.1. Chemicals

Digitoxigenin, digoxigenin, digitoxin, lanatoside C, *β*-acetyldigoxin, and HPLC-grade methanol, were purchased from the Millipore Sigma (St. Louis, Missouri, United States). Digoxin was obtained from the Acros Organics (New Jersey, United States) and lanatoside B was acquired from the Alkemist Labs (Garden Grove, California, United States). LC/MS grade mobile phase A (0.1 % formic acid in water) and B (0.1 % formic acid in acetonitrile) were purchased from Honeywell International Inc (Muskegon, Michigan, United States).

### 2.2. Plant materials

Seeds of *Digitalis lanata Ehrh*. and *Digitalis purpurea L.* were purchased from Strictly Medicinal (Williams, Oregon, USA). The seeds were germinated and maintained in a growth chamber (Invitrogen, Clayton, Missouri, USA) under a 16-hour light period at 25 °C with a relative humidity of 60-80 %. The plants were grown in a soil mix (50 L loosened peat, 25 L perlite, 25 L coarse vermiculite, 57 g triple superphosphate, 57 g bone meal, 85 g calcium hydroxide, 99 g calcium carbonate and 369 g Osmocote 14-14-14). The pair of the first true leaves were collected 5 weeks after germination, immediately frozen in liquid nitrogen and stored in 80 °C before extraction.

### 2.3. Preparation of plant extract

Biological triplicates were prepared from the first pair of true leaves of individual plants grown in separate pots. Leaves of *D. lanata* and *D. purpurea* were freeze-dried for 24 hours using a Labconco FreeZone 2.5 lyophilizer (Kansas, Missouri, USA). The samples were then homogenized using polypropylene pellet pestles (DWK life sciences, NJ, USA) in 1.5 ml tubes. 100 μL of 80% methanol was used to resuspend per mg of dry tissue at room temperature. The samples were vortexed and incubated at 65 °C for 10 min. The extract was then centrifuged at 18,000 g for 10 minutes and the supernatant was filtered through a 0.45 μm multiscreen filter plate (Merck Millipore, Carrigtwohill, Ireland). The samples were stored at - 20 °C before LC/MS analysis.

### 2.4. LC/MS Analysis for cardenolide standards and primary cardenolides in Digitalis leaves

Analysis of cardiac glycosides in *Digitalis* leaf extracts was carried out using a Thermo Scientific Q-Exactive Focus™ with Thermo Scientific UltiMate 3000 UHPLC™, operated in the positive ion mode for electrospray ionization (+ESI). Chromatographic separation was obtained using a Waters XSelect CSH™ C18 HPLC column (3.5 μm particle size, 2.1 mm i.d., 150 mm length) with a mobile phase consisting of water with 0.1 % formic acid (mobile phase A) and acetonitrile with 0.1 % formic acid (mobile phase B). Gradient elution was carried out at a flow rate of 200 μL min^−1^ starting with 10 % mobile phase B followed by a 12-min linear gradient to 95% mobile phase B, held for 2 min, and then brought back to initial conditions in 1 min and equilibrated for 3 min before next injection. The sample injection volume was 10 μL. A full-scan with data-dependent MS^2^ (full MS-ddMS^2^) was used with an inclusion list of the precursor ion masses for target analytes. Settings for full MS: scan range 100-1200m/z, resolution 70,000, automatic gain control (AGC) target of 1E^6^ with a maximum injection time set to automatic. For ddMS^2^, the settings: resolution of 17,500, isolation window of 4.0 m/z; collision energies at 10, 30 and 60 V, AGC target of 1E^5^ and maximum injection time also set to automatic. Quantitative analysis was completed using the vendor-supplied Xcalibur ™ software employing external calibration.

### 2.5. Quantification of cardenolide standards

Individual cardenolide stock solutions at 100 μg g^−1^ were prepared in 100 % methanol and stored at -80 °C. Dilutions of the mixed cardenolides standards were prepared in 80% methanol with concentrations of 0.8 ng g^−1^, 4 ng g^−1^, 20 ng g^−1^, 100 ng g^−1^, 500 ng g^−1^. Cardenolide standards were quantified using the LC/MS method developed above with five technical replicates. Quantitative data processing was performed using the Xcalibur™ software. [M+H]^+^ adducts were used for calibration curves of all cardenolides exception for digitoxin whose [M+Na]^+^ adduct ionized 10-fold better than the [M+H]^+^. The limit of detection (LOD) was defined as 3.3 folds of the standard deviation of the lowest response (*δ)* divided by the slope of the linear calibration curve (*S*), or LOD = 3.3 *δ*/*S*. The limit of quantification (LOQ) was calculated as LOQ = 10 *δ*/*S*.

### 2.6. HRLC/MS analysis to identify other cardenolides in Digitalis leaf extract

To identify other cardenolides that might be present in the leaf samples, the chromatographic method described above was extended to a 60–min method starting at 10 % mobile phase B followed by a 5-min linear gradient to 30 % mobile phase B, followed by a 15-min linear gradient to 70 % mobile phase B, held for 10 mins, followed by a 10-min linear gradient to 95% mobile phase B, also held for 10 min. After which, the gradient was returned to initial conditions within 2 min and equilibrated for 8 mins before the next injection. A similar full MS-ddMS^2^ was performed with an inclusion list of all the cardenolides that could potentially be present in the samples. For less abundant cardenolide that did not fragment in the full MS-ddMS^2^ method, a selected ion monitoring with MS^2^ (SIM-ddMS^2^) method using the 60-min chromatographic condition with an inclusion list of identified precursor ions was developed to obtain MS^2^ spectra. Settings for the SIM method followed: the resolution of 35,000, AGC target of 5E4 with a maximum injection time set to automatic and a ddMS^2^ resolution of 17,500, isolation window of 4.0 m/z; collision energies at 10, 30 and 60 V, maximum injection time set to automatic. Data processing involved extracting the precursor ions of the suspect cardenolides and their MS^2^ spectra in Xcalibur™ with mass error set to 5 ppm and retention time match. Semi-quantification of identified cardenolides was performed by summing peak areas of both [M+H]^+^ and [M+Na]^+^ adducts. The percentage of total cardenolides was calculated as the ratio of the peak area for individual cardenolide divided by that of total cardenolides, given that cardenolides ionized similarly as determined by seven cardenolide standards.

### 2.7. Prediction of physicochemical properties

The octanol-water partition coefficients of cardenolides was predicted using XLogP3 3.0 software [18]. Numbers of rotatable bonds, hydrogen bond donors and acceptors were calculated by Cactvs 3.4.5.11 version [19]. The theoretical exact masses were computed using the Compass Isotope Pattern calculator (Bruker, Bermen, Germany).

## 3. Results and Discussion

### 3.1. The MS spectra of cardiac glycosides reveal structures of both the aglycones and the sugar units

To study how cardiac glycosides fragment during CID, available pure standards of cardenolides were first analyzed using a Thermo Scientific Q-Exactive Focus™ liquid-chromatography tandem mass spectrometry system (QE) with electrospray ionization and a high-resolution Orbitrap™ analyzer, in the positive ion mode. Both the [M+H]^+^ and the [M+Na]^+^ adducts were identified within Δ m/z less than 5 ppm but only the [M+H]^+^ ions generated subsequent product ions with a comprehensive fragmentation profile (Fig. 2). Digoxin was broken down through the sequential loss of the three digitoxose units (131.0703 m/z). The resulting monosaccharide cation was resonance stabilized between the hemiacetal oxygen and the anomeric carbon (Fig. 2A, structure G). The digoxigenin aglycone underwent a three-step dehydration (18.0106 m/z) characteristic for hydroxyl groups. The signature ions for digoxigenin were: 391.2479 m/z, 373. 2373 m/z, 355.2268 m/z, 337.2162 m/z, resulting from stepwise dehydration. As expected, these aglycone peaks were also observed in *β*-acetyldigoxin, lanatoside C, and digoxigenin, which all contain the same aglycone (Fig. 4A, Supplemental Fig. 1). A two-step dehydration was observed with digitoxin since it only has two hydroxyl groups. The signature ions for aglycone digitoxigenin were: 375.2530 m/z, 357.2424 m/z, 339,2319 m/z, which were also detected in digitoxigenin. In *β*-acetyldigoxin, the additional acetyldigitoxose unit (173.0808 m/z) was detected [20] (Supplemental Fig. 1). All identified product ions were with minimum Δ m/z compared to the theoretical exact masses. The observation of sequential sugar loss and dehydration of cardenolides agreed with previous reports but with higher mass resolution, allowing unambiguous assignment of precursor and product ions [8,21].

**Fig. 2.**
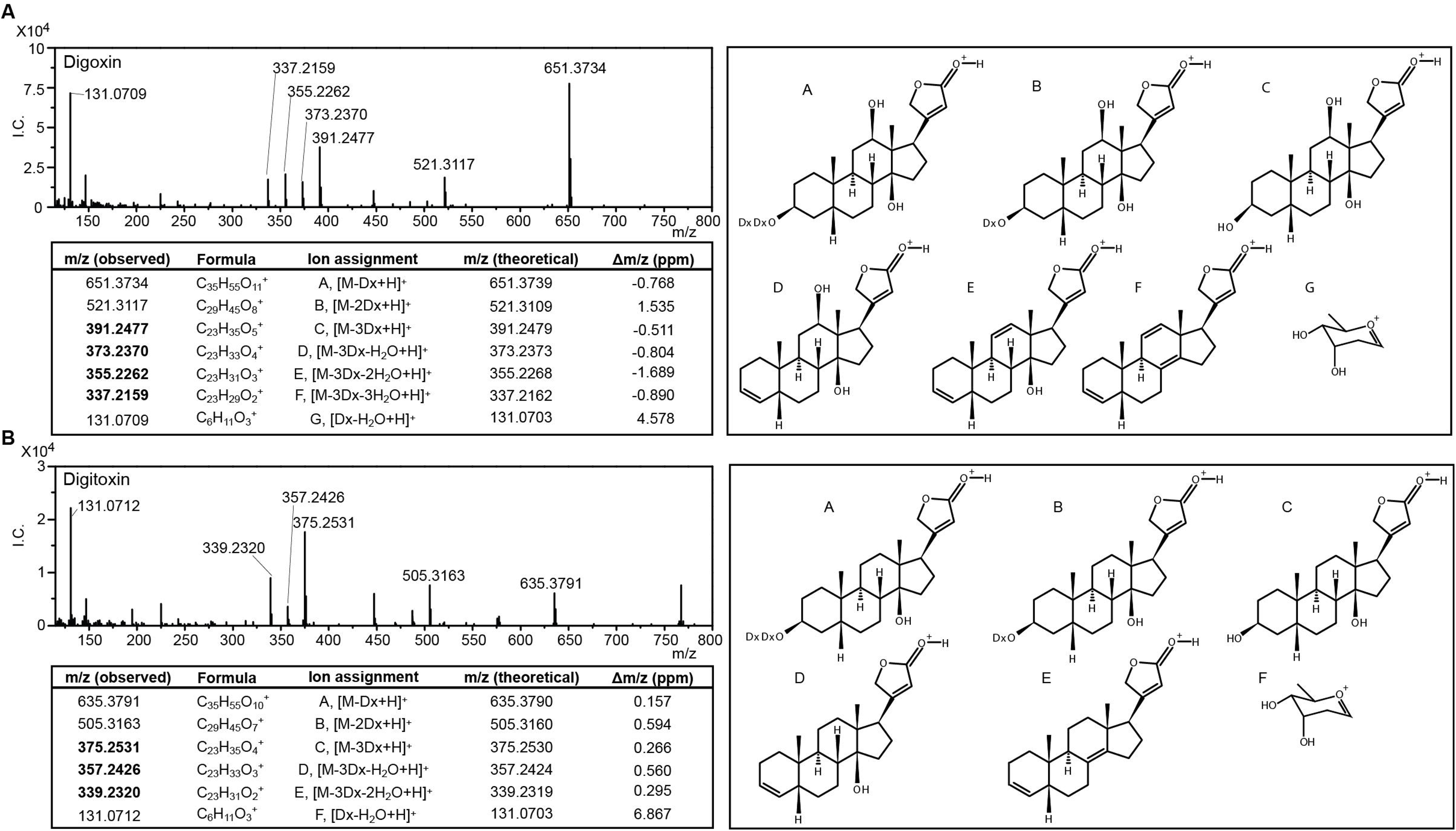
MS^2^ product ion mass spectra from [M+H]^+^ adducts of digoxin (A) and digitoxin (B) standards with putative ion structures. Signature product ions of digoxigenin and digitoxigenin aglycones are highlighted. I.C.: ion counts; ppm: parts per million; Dx: digitoxose unit

**Fig. 3.**
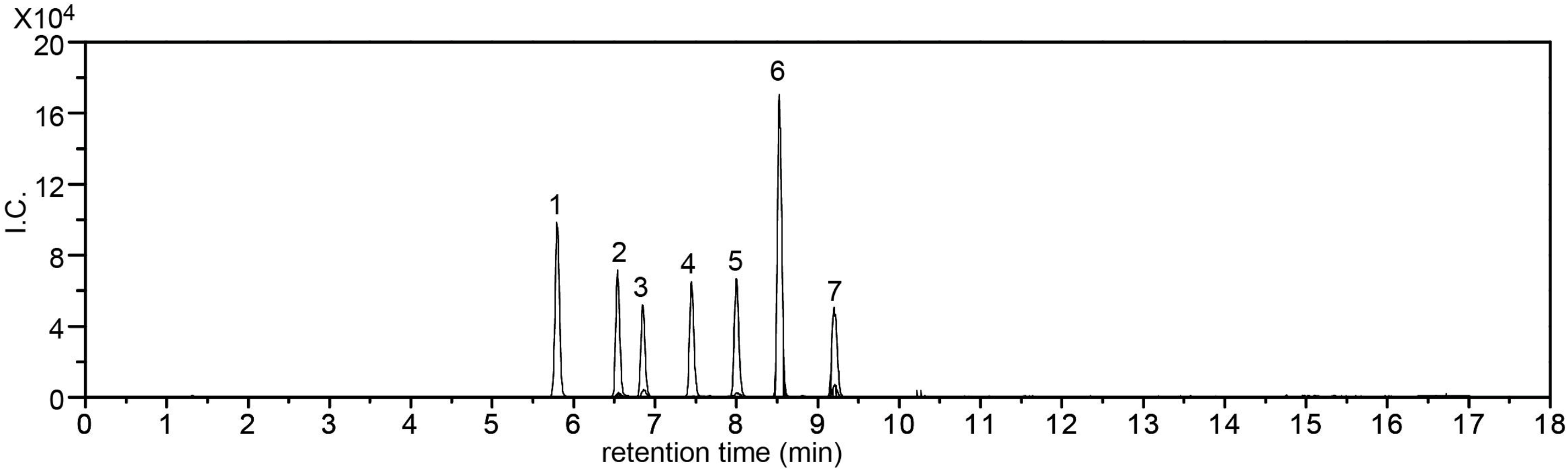
Overlaid full-scan chromatograms of cardiac glycoside standards. Chromatograms of [M+H]^+^ are shown for all pure compounds except digitoxin (peak 7) for which the [M+Na]^+^ adduct is shown. The concentrations for each compound are 500 ng g^−1^ except digoxigenin and digitoxigenin for which 100 ng g^−1^ is used. The peaks are: 1, digoxigenin; 2, lanatoside C; 3, digoxin; 4, lanatoside B; 5, *β*-acetyldigoxin; 6, digitoxigenin; 7, digitoxin.

**Fig. 4.**
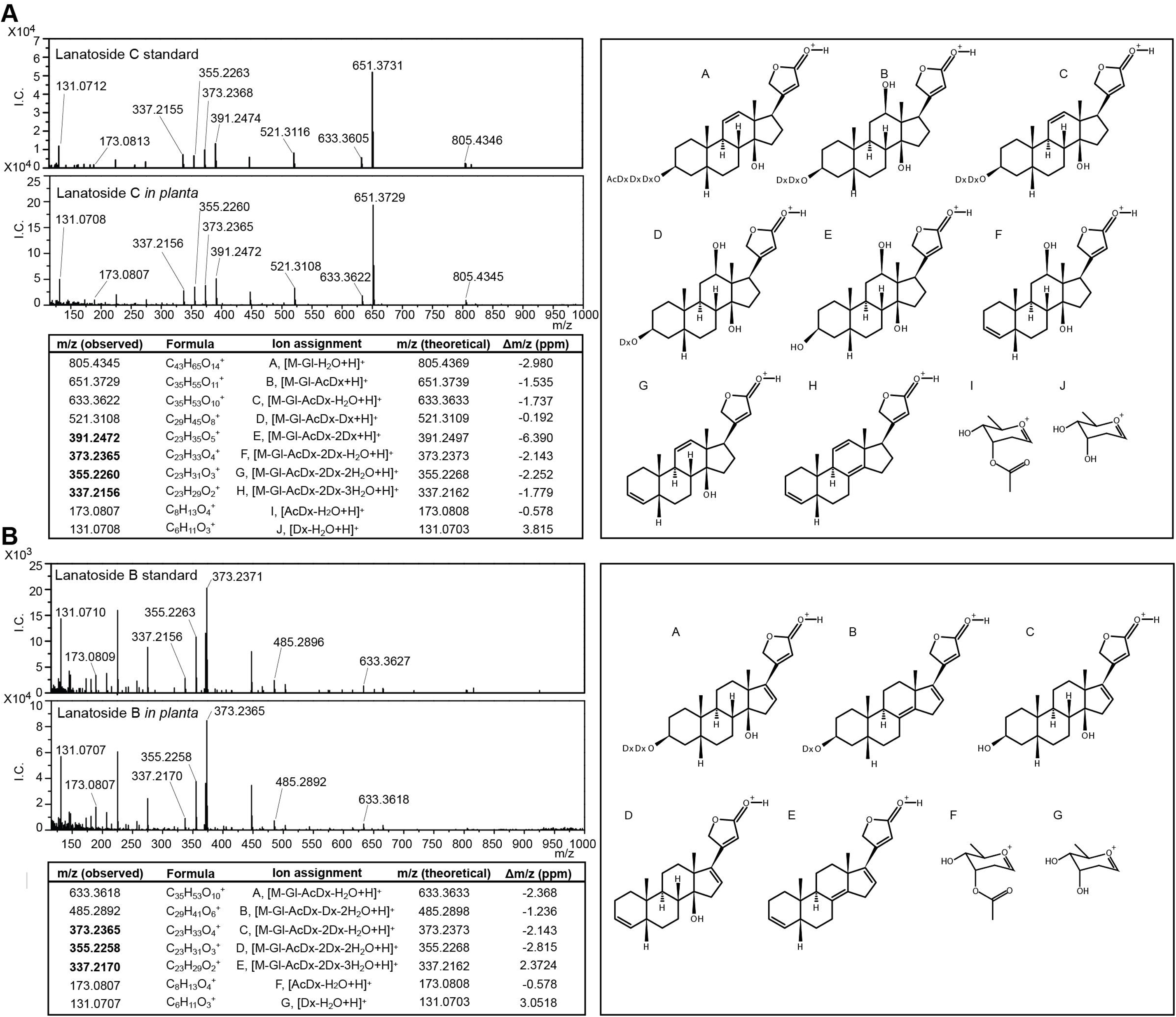
Comparison of MS^2^ spectra of lanatoside C and lanatoside B [M+H]^+^ adducts *in planta* versus pure standards. (A) MS^2^ spectra of lanatoside C and putative ion structures; (B) MS^2^ spectra of lanatoside B and putative ion structures. Signature product ions of digoxigenin and gitoxigenin aglycones are highlighted. I.C.: ion counts; ppm: parts per million; Gl: glucose unit; Dx: digitoxose unit; AcDx: acetyldigitoxose unit.

### 3.2. Quantification of cardiac glycosides by high-resolution MS (HRMS)

Quantification of authentic standards of cardenolides was achieved in methanol using a 17-minute method with a reverse-phase C18 column followed by Orbitrap™ detection. The sequence of elution for cardenolides with identical sugar units but different aglycones followed digoxigenin → gitoxigenin → digitoxigenin, which agrees with previous HPLC and LC/MS reports (Fig. 3 and Table 1) [7,8]. Digitoxigenin with two hydroxyl groups is less polar than digoxigenin or gitoxigenin with three hydroxyl groups, therefore eluted later. Lanatoside C and lanatoside B are regioisomers that differ only in the position of one hydroxyl group (12-OH v.s. 16-OH). Lanatoside B eluted after lanatoside C because the 16-OH of lanatoside B is sterically hindered by the 17- *γ*-butyrolactone ring, rendering it less polar than lanatoside C. Indeed, the log P value of lanatoside B was predicted to be larger than that of lanatoside C (Fig. 1). The elution of cardenolides with the same aglycone but different sugar units followed: aglycone →primary cardenolides →secondary cardenolides →acetylated secondary cardenolides, as exemplified by digoxigenin, lanatoside C, digoxin, and *β*-acetyldigoxin (Fig. 3). The [M+H]^+^ precursor ions were quantified because of broader linear ranges and lower LODs than [M+Na] ^+^ except for digitoxin whose sodium adduct ionized 10-fold better than the proton adduct. The calibration curves obtained for *β*-acetyldigoxin, digitoxigenin, and digoxigenin have a linear range of 0.8 -500 ng g^−1^ with limits of detection (LOD) that vary from 0.09 to 8.3 ng g^−1^. The linear ranges obtained were about 10-fold broader than previous LC/MS methods except for digitoxin [8,12]. The LODs were comparable to earlier reports or lower as of *β*-acetyldigoxin [8]. With a sensitive method in hand, we next examined cardenolides in crude *Digitalis* extracts.

**Table 1.** Quantification of cardiac glycoside standards. LOD, limit of detection; LOQ, limit of quantification, R^2^: correlation coefficient. Data were obtained from five injections.

| Compound | Retention time (min) | Ion adduct | Slope | Intercept | Linear Range (ng g<sup>-1</sup>) | $R^2$ | LOD (ng g<sup>-1</sup>) | LOQ (ng g<sup>-1</sup>) |
| --- | --- | --- | --- | --- | --- | --- | --- | --- |
| Digoxigenin | 5.81 | [M+H]<sup>+</sup> | 310513 | 1000000 | 0.8-500 | 0.9993 | <0.45 | <1.35 |
| Lanatoside C | 6.56 | [M+H]<sup>+</sup> | 47918 | 133358 | 4-500 | 0.9996 | <2.71 | <8.21 |
| Digoxin | 6.84 | [M+H]<sup>+</sup> | 36694 | 134761 | 4-500 | 0.9995 | <2.74 | <8.30 |
| Lanatoside B | 7.48 | [M+H]<sup>+</sup> | 199836 | 23071 | 4-500 | 0.9991 | <1.79 | <5.44 |
| β-Acetyldigoxin | 8.02 | [M+H]<sup>+</sup> | 50710 | 133055 | 0.8-500 | 0.9994 | <0.09 | <0.27 |
| Digitoxigenin | 8.51 | [M+H]<sup>+</sup> | 573174 | 1000000 | 0.8-500 | 0.9997 | <0.34 | <1.12 |
| Digitoxin | 9.22 | [M+Na]<sup>+</sup> | 140984 | 24048 | 4-100 | 0.9999 | <3.00 | <7.82 |

### 3.3. The MS spectra of primary cardenolides in planta

The same 17-minute LC/MS^2^ method was used to analyze cardenolides in the fresh-leave extract of *D. lanata*. Lanatoside C standard had near-identical fragmentation patterns as digoxin but with two additional fragments corresponding to the acetyldigoxin unit (805.4369 m/z) and the acetyldigitoxose unit (173.808 m/z) respectively (Fig. 4A). Interestingly, lanatoside B, a regioisomer of lanatoside C, had all aglycone signature peaks of lanatoside C but missing the 391.2497 m/z peak, the protonated gitoxigenin without any dehydration (Fig. 4B). All fragments of lanatoside B with the gitoxigenin aglycone underwent at least one dehydration, indicating the 16-OH group was readily eliminated during ionization. This can be understood that the resulting cation of 16-OH dehydration is stabilized by both the extended conjugate π bounds as well as the formation of a trisubstituted double bond. Thus, the presence or absence of the 391.2497 m/z peak distinguished the digoxigenin from the gitoxigenin aglycone. The [M+H]^+^ precursor ions of lanatoside B and C detected in *D. lanata* extracts had identical retention times and fragmentation patterns as the pure standards, verifying the presence of both primary cardenolides in *D. lanata*.

Other primary cardenolides, lanatoside A, D, and E were also identified and structurally annotated in *D. lanata* leaf extracts (Fig. 5). Single elution peak was shown for all [M+H]^+^ precursor ions of lanatoside A, D, and E within 2 ppm window of the theoretical exact mass. The primary cardenolides followed the elution sequence: lanatoside D → lanatoside C → lanatoside B → lanatoside E → lanatoside A, corresponding well with their log P values (Fig. 1). Lanatoside D with hydroxyl groups on both 12- and 16-position is the most polar in the family, therefore eluted first. Although isomers of each other, lanatoside C eluted before B because the 12-OH encountered less steric hindrance than the 16-OH. Lanatoside E has a formate ester instead of a hydroxyl group at the 16^-^ position rendering it less polar than lanatoside B, thus eluted afterward. Lanatoside A is the most non-polar compound due to the lack of hydroxylation at either 12- or 16- position and was the last to elute. The observed elution sequence from a C18 column concurred with previous reports [7].

**Fig. 5.**
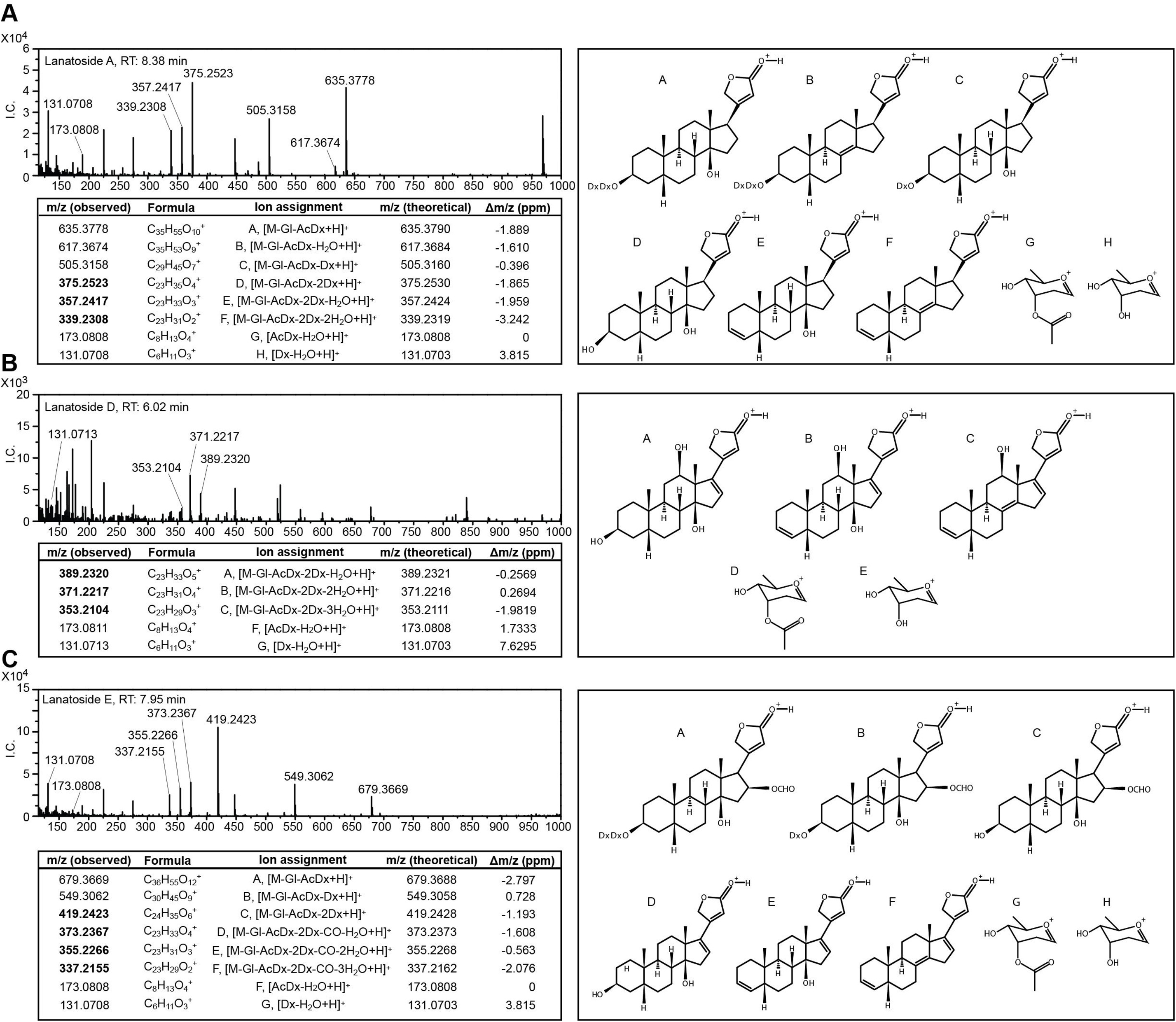
MS^2^ product ion mass spectra from [M+H]^+^ adduct of lanatoside A, D, and E with putative ion structures. Signature product ions of digitoxigenin, diginatigenin, and gitaloxigenin aglycones are highlighted. I.C.: ion counts; ppm: parts per million; Gl: glucose unit; Dx: digitoxose unit; AcDx: acetyldigitoxose unit.

The MS^2^ spectra of lanatoside A, D, and E also showed stepwise loss of sugar units and dehydration of the hydroxyl groups of the aglycone (Fig. 5). Just like lanatoside B and C, the 173.0808 m/z peak and the 131.0705 m/z peak corresponding to the acetyldigitoxose and digitoxose moieties were identified in lanatoside A, D, and E. As expected, lanatoside A fragmentations included the aglycone signatures identical to those in digitoxigenin and digitoxin (375.2530 m/z, 357.2424 m/z, 339,2319 m/z) with 18.0106 m/z intervals (Fig. 5A). Lanatoside D with hydroxylation at both the 12- and the 16- position had unique aglycone signatures (389.2321 m/z, 371.2216 m/z, 353.2111 m/z) that were 15.9948 m/z larger (the additional oxygen) than the corresponding aglycone peaks of lanatoside B or C (373. 2373 m/z, 355.2268 m/z, 337.2162 m/z) (Fig. 5B). Just like lanatoside B, the 16-OH in lanatoside D was easily eliminated during ESI due to the resulting resonance stabilized cation. It was also found that lanatoside D was in negligible amount in *D. lanata* which disagreed with previous work using HPLC [7]. It was unlikely that lanatoside D ionize poorly in our assay because close analogs such as lanatoside B or C ionized well. The higher abundance of lanatoside D detected by HPLC before could result from the unresolved co-eluents from the plant mixture. It was also possible that the *D. lanata* plant we grew produced less lanatoside D than the plant used by other groups since *D. lanata* was an outbreeding species with genetic variances. The spectra of lanatoside E showed a unique 419.2428 m/z peak corresponding to the gitaloxigenin aglycone and other dehydrated aglycone peaks (373. 2373 m/z, 355.2268 m/z, 337.2162 m/z) identical to those in lanatoside B and C (Fig 5C). The decrease in mass from 419.2428 m/z to the 373.2373 m/z corresponded to the elimination of a formic acid (46.0055 m/z) generating the same resonance stabilized cation found in the spectra of lanatoside B. Thus far, all aglycones were distinguished based upon their MS/MS spectra, we next examined all other possible cardenolides in two most commonly studied *Digitalis* species.

### 3.4. Identification and semi-quantification of cardenolides in D. lanata and D. purpurea leaf extracts

To achieve a better separation, a 60-minute LC method was developed for the analysis of other cardenolides in *Digitalis*. Fresh leaves were freeze-dried and extracted in 80% methanol to avoid partial hydrolysis of primary cardenolides. Overall, a total of 17 and 7 cardenolides were detected in *D. lanata* and *D. purpurea* respectively (Fig. 6). The cardenolide profiles were unique and distinct in these two closely related species, which concurred with previous reports [7,22]. Overall, *D. lanata* had a higher concentration and more varieties of cardenolides compared with *D. purpurea*, consistent with the usage of the former for producing digoxin and the latter as an ornamental plant.

**Fig. 6.**
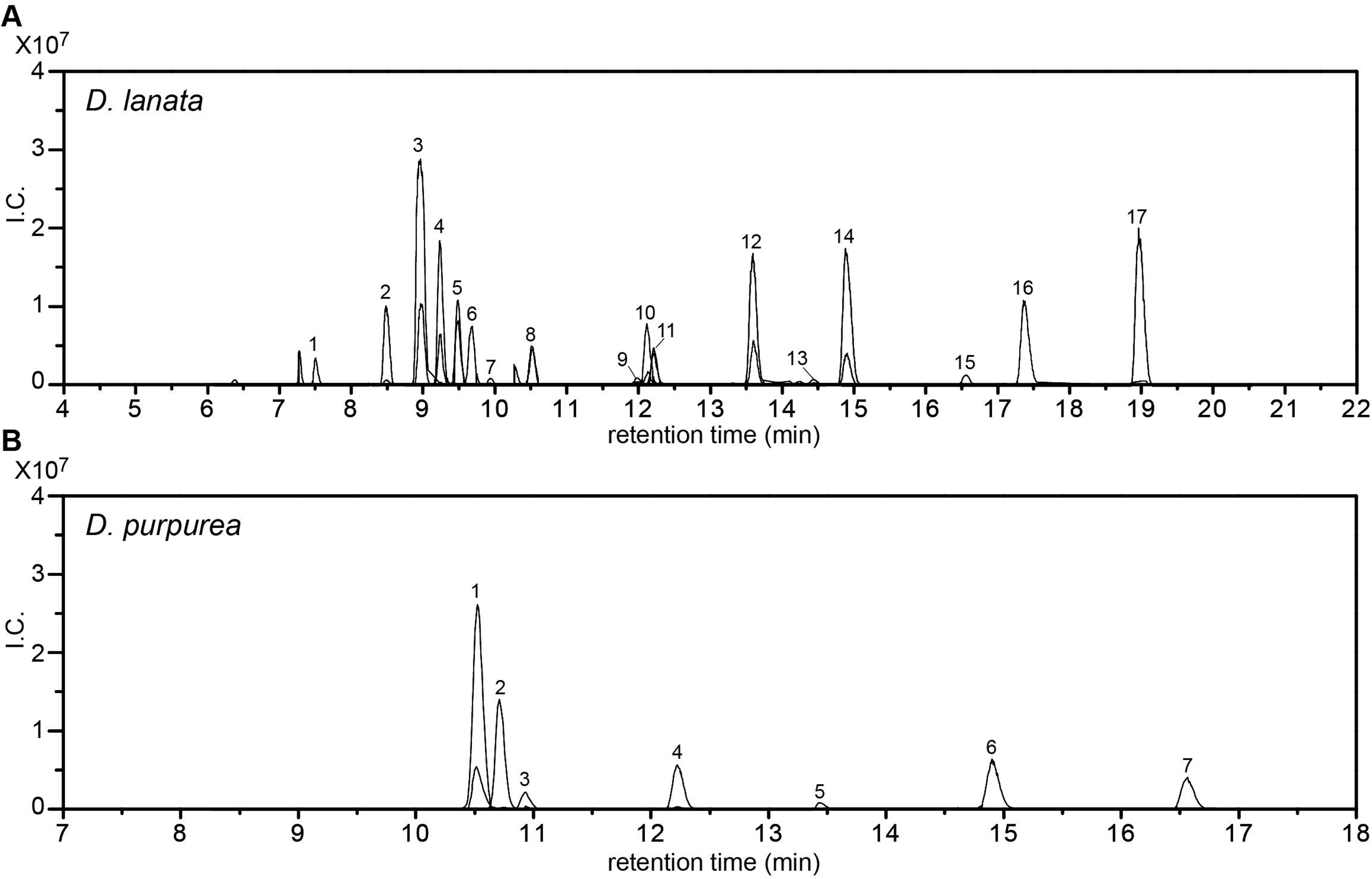
Overlaid SIM chromatograms of cardenolides detected in leaves of *D. lanata* and *D. purpurea*. Both [M+H]^+^ and [M+Na]^+^ adducts are shown for each compound. (A) Cardenolides identified in *D. lanata.* The peaks are: 1, digitalinum verum; 2, glucoverodoxin; 3, glucodigifucoside; 4, odorobioside G; 5, lanatoside C; 6, glucolanadoxin; 7,digoxin; 8, verodoxin; 9, Abbreviated name: Gl-Dx-Dx-A; 10, α-acetyldigoxin; 11, lanatoside B; 12, lanatoside E; 13, digitoxin bis-digitoxide; 14, lanatoside A; 15, digitoxin; 16, *α/β*-acetylgitaloxin; 17,, *α/β*-acetyldigitoxin. (B) Cardenolides identified in *D. purpurea*. The peaks are:1, verodoxin; 2, digitoxigenin fucoside; 3, purpurea glycoside B; 4, glucogitaloxin; 5, purpurea glycoside A; 6, gitaloxin; 7, digitoxin.

In particular, purpurea glycosides were not identified in *D. lanata*, and lanatosides were not detected in *D. purpurea.* Additionally, acetylated cardenolides and cardenolides with digoxigenin and diginatigenin aglycones were only found in *D. lanata*, which strongly indicates that the D-digitoxose 3-acetyltransferase and the 12- steroid hydroxylase are only present in *D. lanata* but not in *D. purpurea*. The chemical formula of all [M+H]^+^ precursor and product ions were unambiguously determined by HRMS with minimum Δ m/z (Tables 2 and 3). The MS/MS spectra of each identified cardenolide contained signature peaks for both the aglycone and the sugar units[20] (Supplemental Fig. 2). The elution followed the C→B→E→A sequence for cardenolides with the same sugar units but different aglycones just like authentic standards. Although lanatoside B and C are constitutional isomers with the same exact masses, they were distinguished by the retention times identical to the respective pure compounds and the presence or absence of the 391.2497 m/z peak in the MS/MS spectra. The *α*- isomer of acetyldigoxin was assigned because its retention time was different from the pure compound *β*-acetyldigoxin. The exact diastereomers of acetylgitaloxin and acetyldigitoxin were not distinguishable due to the lack of authentic standards. In future, NMR will be necessary to determine the stereochemistry of these two compounds.

**Table 2.**
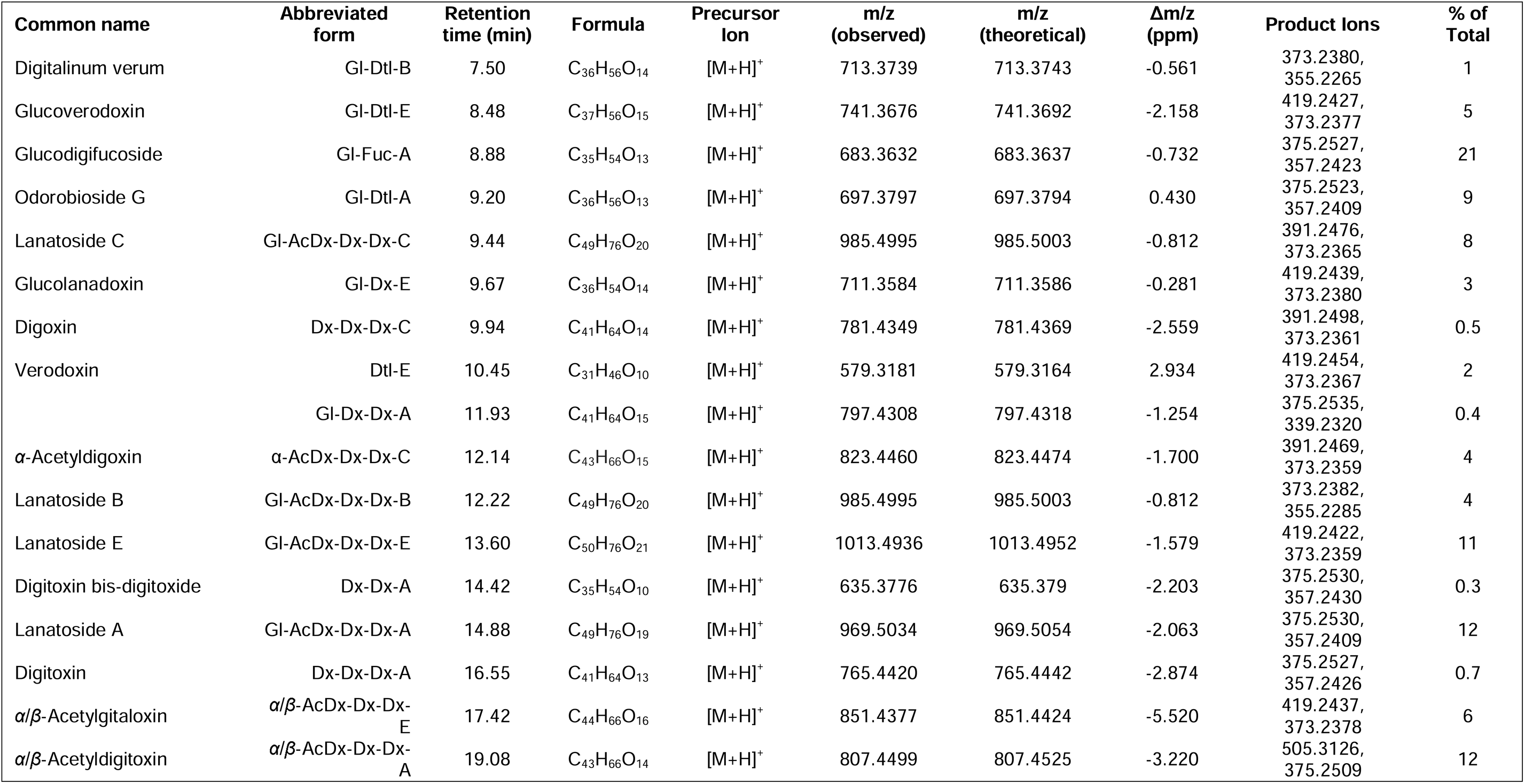
MS^2^ dectection of cardenolides from *Digitalis lanata* fresh leaves. Aglycones: A, digitoxigenin; B, gitoxigenin; C, digoxigenin; D, diginatigenin; E, gitaloxigenin. Sugars: AcDx, acetyldigitoxose; Dx, digitoxose; Gl, glucose; Dtl, digitalose; Fuc, fucose. The relative abundance of each cardenolide is calculated as % of total using both [M+H]^+^ and [M+Na]^+^ adducts. Cardenolides identified in at least two out of three biological replicates are listed.

| Common name | Abbreviated form | Retention time (min) | Formula | Precursor Ion | m/z (observed) | m/z (theoretical) | Δm/z (ppm) | Product Ions | % of Total |
| --- | --- | --- | --- | --- | --- | --- | --- | --- | --- |
| Digitalinum verum | Gl-Dtl-B | 7.50 | C<sub>36</sub>H<sub>56</sub>O<sub>14</sub> | [M+H]<sup>+</sup> | 713.3739 | 713.3743 | -0.561 | 373.2380, 355.2265 | 1 |
| Glucoverodoxin | Gl-Dtl-E | 8.48 | C<sub>37</sub>H<sub>56</sub>O<sub>15</sub> | [M+H]<sup>+</sup> | 741.3676 | 741.3692 | -2.158 | 419.2427, 373.2377 | 5 |
| Glucodigifucoside | Gl-Fuc-A | 8.88 | C<sub>35</sub>H<sub>54</sub>O<sub>13</sub> | [M+H]<sup>+</sup> | 683.3632 | 683.3637 | -0.732 | 375.2527, 357.2423 | 21 |
| Odorobioside G | Gl-Dtl-A | 9.20 | C<sub>36</sub>H<sub>56</sub>O<sub>13</sub> | [M+H]<sup>+</sup> | 697.3797 | 697.3794 | 0.430 | 375.2523, 357.2409 | 9 |
| Lanatoside C | Gl-AcDx-Dx-Dx-C | 9.44 | C<sub>49</sub>H<sub>76</sub>O<sub>20</sub> | [M+H]<sup>+</sup> | 985.4995 | 985.5003 | -0.812 | 391.2476, 373.2365 | 8 |
| Glucolanadoxin | Gl-Dx-E | 9.67 | C<sub>36</sub>H<sub>54</sub>O<sub>14</sub> | [M+H]<sup>+</sup> | 711.3584 | 711.3586 | -0.281 | 419.2439, 373.2380 | 3 |
| Digoxin | Dx-Dx-Dx-C | 9.94 | C<sub>41</sub>H<sub>64</sub>O<sub>14</sub> | [M+H]<sup>+</sup> | 781.4349 | 781.4369 | -2.559 | 391.2498, 373.2361 | 0.5 |
| Verodoxin | Dtl-E | 10.45 | C<sub>31</sub>H<sub>46</sub>O<sub>10</sub> | [M+H]<sup>+</sup> | 579.3181 | 579.3164 | 2.934 | 419.2454, 373.2367 | 2 |
|  | Gl-Dx-Dx-A | 11.93 | C<sub>41</sub>H<sub>64</sub>O<sub>15</sub> | [M+H]<sup>+</sup> | 797.4308 | 797.4318 | -1.254 | 375.2535, 339.2320 | 0.4 |
| α-Acetyldigoxin | α-AcDx-Dx-Dx-C | 12.14 | C<sub>43</sub>H<sub>66</sub>O<sub>15</sub> | [M+H]<sup>+</sup> | 823.4460 | 823.4474 | -1.700 | 391.2469, 373.2359 | 4 |
| Lanatoside B | Gl-AcDx-Dx-Dx-B | 12.22 | C<sub>49</sub>H<sub>76</sub>O<sub>20</sub> | [M+H]<sup>+</sup> | 985.4995 | 985.5003 | -0.812 | 373.2382, 355.2285 | 4 |
| Lanatoside E | Gl-AcDx-Dx-Dx-E | 13.60 | C<sub>50</sub>H<sub>76</sub>O<sub>21</sub> | [M+H]<sup>+</sup> | 1013.4936 | 1013.4952 | -1.579 | 419.2422, 373.2359 | 11 |
| Digitoxin bis-digitoxide | Dx-Dx-A | 14.42 | C<sub>35</sub>H<sub>54</sub>O<sub>10</sub> | [M+H]<sup>+</sup> | 635.3776 | 635.379 | -2.203 | 375.2530, 357.2430 | 0.3 |
| Lanatoside A | Gl-AcDx-Dx-Dx-A | 14.88 | C<sub>49</sub>H<sub>76</sub>O<sub>19</sub> | [M+H]<sup>+</sup> | 969.5034 | 969.5054 | -2.063 | 375.2530, 357.2409 | 12 |
| Digitoxin | Dx-Dx-Dx-A | 16.55 | C<sub>41</sub>H<sub>64</sub>O<sub>13</sub> | [M+H]<sup>+</sup> | 765.4420 | 765.4442 | -2.874 | 375.2527, 357.2426 | 0.7 |
| α/β-Acetylgitaloxin | α/β-AcDx-Dx-Dx-E | 17.42 | C<sub>44</sub>H<sub>66</sub>O<sub>16</sub> | [M+H]<sup>+</sup> | 851.4377 | 851.4424 | -5.520 | 419.2437, 373.2378 | 6 |
| α/β-Acetyldigitoxin | α/β-AcDx-Dx-Dx-A | 19.08 | C<sub>43</sub>H<sub>66</sub>O<sub>14</sub> | [M+H]<sup>+</sup> | 807.4499 | 807.4525 | -3.220 | 505.3126, 375.2509 | 12 |

**Table 3.** MS^2^ detection of cardenolides from *Digitalis purpurea* fresh leaves. Aglycones: A, digitoxigenin; B, gitoxigenin; E, gitaloxigenin. Sugars: AcDx, acetyldigitoxose; Dx, digitoxose; Gl, glucose; Dtl, digitalose; Fuc, fucose. The relative abundance of each cardenolide is calculated as % of total using both [M+H]^+^ and [M+Na]^+^ adducts. Cardenolides identified in at least two out of three biological replicates are listed.

| Common name | Abbreviated form | Retention time (min) | Formula | Precursor Ion | m/z (observed) | m/z (theoretical) | $\Delta m/z$ (ppm) | Product Ions | % of Total |
| --- | --- | --- | --- | --- | --- | --- | --- | --- | --- |
| Verodoxin | Dtl-E | 10.51 | C<sub>31</sub>H<sub>46</sub>O<sub>10</sub> | [M+H]<sup>+</sup> | 579.3160 | 579.3164 | -0.690 | 419.2422, 373.2368 | 48 |
| Digitoxigenin fucoside | Fuc-A | 10.69 | C<sub>29</sub>H<sub>44</sub>O<sub>8</sub> | [M+H]<sup>+</sup> | 521.3108 | 521.3109 | -0.192 | 375.2533, 357.2417 | 20 |
| Purpurea glycoside B | Gl-Dx-Dx-Dx-B | 10.96 | C<sub>47</sub>H<sub>74</sub>O<sub>19</sub> | [M+H]<sup>+</sup> | 943.4911 | 943.4897 | 1.484 | 373.2370, 355.2273 | 3 |
| glucogitaloxin | Gl-Dx-Dx-Dx-E | 12.20 | C<sub>48</sub>H<sub>74</sub>O<sub>20</sub> | [M+H]<sup>+</sup> | 971.4819 | 971.4846 | -2.779 | 419.2421, 355.2257 | 9 |
| Purpurea glycoside A | Gl-Dx-Dx-Dx-A | 13.44 | C<sub>47</sub>H<sub>74</sub>O<sub>18</sub> | [M+H]<sup>+</sup> | 927.4907 | 927.4948 | -4.421 | 375.2421, 339.2339 | 1 |
| Gitaloxin | Dx-Dx-Dx-E | 14.89 | C<sub>42</sub>H<sub>64</sub>O<sub>15</sub> | [M+H]<sup>+</sup> | 809.4303 | 809.4318 | -1.853 | 419.2424, 373.2370 | 11 |
| Digitoxin | Dx-Dx-Dx-A | 16.55 | C<sub>41</sub>H<sub>64</sub>O<sub>13</sub> | [M+H]<sup>+</sup> | 765.4406 | 765.4420 | -1.829 | 375.2517, 357.2424 | 7 |

Peak areas of precursor ions were quantified to determine the relative abundance of each compound, given that cardenolide standards ionized similarly (Tables 2 and 3). In *D. lanata*, lanatoside A, E, and C were the major primary cardenolides while lanatoside B was a minor constituent, and lanatoside D could not be reliably detected (Table 2). The total fraction of primary cardenolide was ∼34%, less than previously reported [7]. Partial hydrolysis of primary cardenolides during extraction was minimum since the hydrolysis products digitoxin and digoxin were only found in trace amounts. Interestingly, glucodigifucoside (Gl-Fuc-A) constituted ∼21 % of total cardiac glycosides in *D. lanata*. In *D. purpurea*, purpurea glycoside E, B, and A made up 13 % of total cardiac glycoside, which was also less than a previous report [22] (Table 3). Surprisingly, verodoxin and digitoxigenin fucoside accounted for ∼48 % and ∼20 % of total cardenolides. Secondary cardenolides gitaloxin and digitoxin were also higher than those in *D. lanata*. Partial hydrolysis of purpurea glycosides during the extraction process was unlikely because the same extraction method was applied to both species. Nevertheless, pure standards are necessary for future unbiased quantifications since the ion abundance is dependent on how well each cardiac glycoside ionizes.

According to the minimum reporting standards for chemical analysis proposed by the Chemical Analysis Working Group [23-25], cardenolides with pure standards available including lanatoside B and C, digoxin, and digitoxin were classified as level 1 identified compounds, based on identical retention times, accurate masses, and MS/MS spectra. The rest of the cardenolides detected belonged to the level 2b or 3, putatively characterized compounds as the spectra of these compounds are not found in the available spectral libraries such as Metlin and Global Natural Product Social Molecular Networking (GNPS) [26,27]. Nevertheless, the cardenolides observed were more than likely to be correctly annotated because 1, they were all identified in the respective species of *Digitalis* previously and were validated by methods including NMR [[7,22,28-32]; 2, HRMS enabled definite determination of formula for both precursor and product ions; 3, no purpurea glycosides were identified in *D. lanata* and no lanatosides were detected in *D. purpurea*, validating the specificity of the method; 4. the elution sequence of cardenolides on a C18 column highly resembled preceding reports [7]. For *D. lanata*, 17 out of 59 cardenolides previously identified by HPLC were confirmed and structurally annotated in our method. HPLC alone is insufficient to unambiguously detect cardenolides because co-eluents such as flavonoid glycosides may coelute with cardenolide standards, resulting in the previous erroneous identification of purpurea glycosides and oleandrin aglycons in *D. lanata* leaves [7].

## 4. Conclusion

In this work, we have provided accurate masses and fragmentations of cardiac glycosides in leaf extracts of two commonly studies species of *Digitalis* using LC coupled with HRMS. Closely related steroid aglycones (digitoxigenin, gitoxigenin, digoxigenin, diginatigenin, gitaloxigenin) can be distinguished by their unique fragmentation patterns. Signature peaks for sugar units (digitoxose, fucose, glucose, acetyldigitoxose, digitalose) have also been identified. This method is better suited for quantifying cardenolides compared with previous HPLC based methods because of the 6 ∼ 30-fold broader linear ranges and ∼1000-fold lower LODs [33]. 17 and 7 cardenolides have been identified in *D. lanata* and *D. purpurea* respectively. It is worth noticing that many cardiac glycosides remain unidentified in the samples because the compounds were detected with only the aglycone recognizable but the sugar units were difficult to interpret. This is the first time an MS method reported for characterizing cardiac glycosides in crude plant extracts. It paves the way for accurate quantification of cardiac glycosides in different species of *Digitalis*, different batches of plant material, different tissues of the same plant, or plants undergoing various stresses, that ultimately sheds light on the biosynthesis and regulation of this family of pharmaceutically important plant natural products.

## Acknowledgments

We would like to thank Dr. Valerie Frerichs at the Chemistry Instrumentation Center at University at Buffalo for stimulating discussions throughout the experiments and careful editing of the manuscript. This work was funded by the generous support from the SUNY Research Foundation.

