## Supplemental Figures for "Profiling and structural analysis of cardenolides in two species of Digitalis using liquid chromatography coupled with high-resolution mass spectrometry"

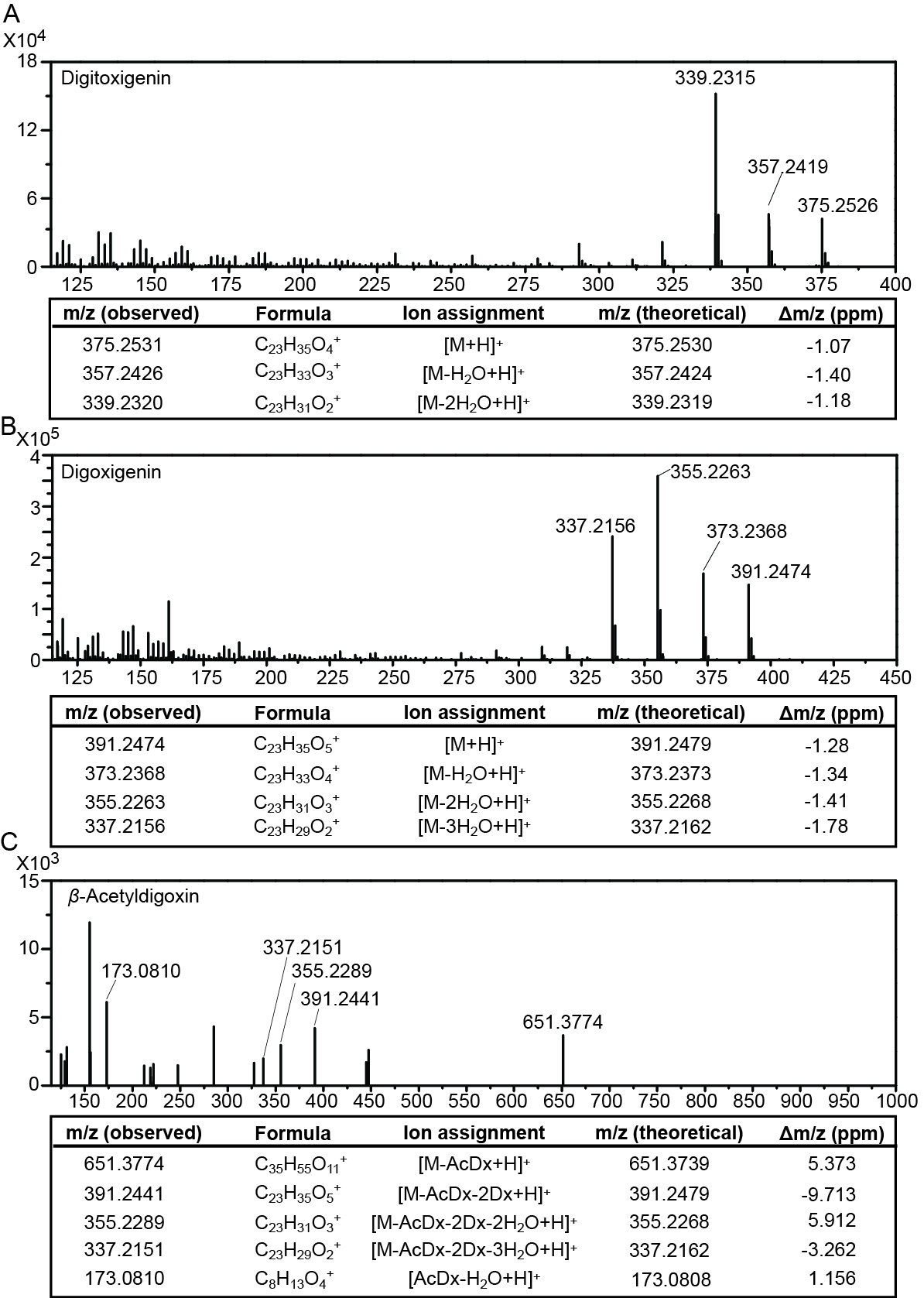

**Fig. S1.** MS^2^ product ion mass spectra from [M+H]^+^ adducts of digitoxigenin (A), digoxigenin (B), and *β*-acetyldigoxin (C) standards with putative ion structures. I.C.: ion counts; ppm: parts per million; Dx: digitoxose unit.

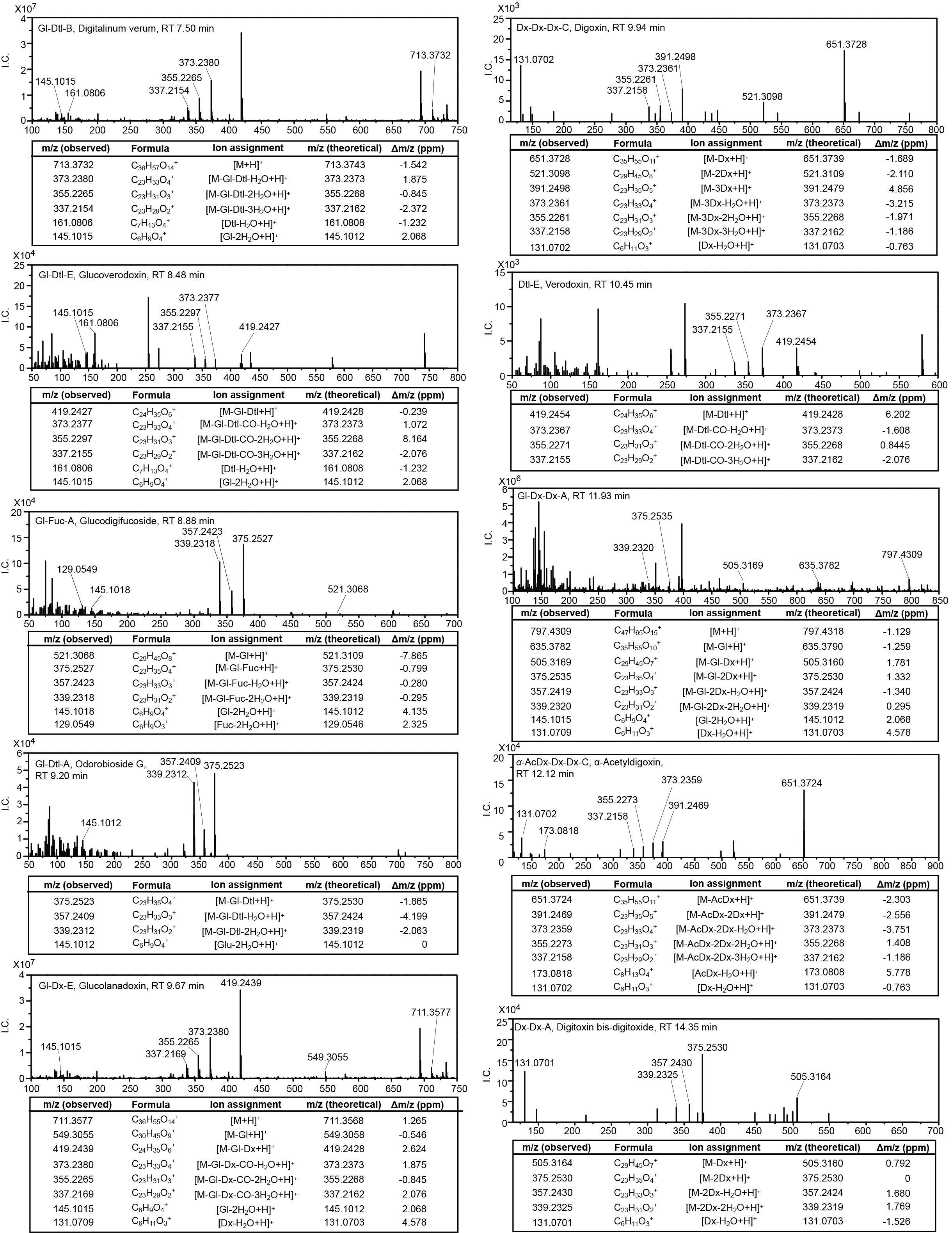

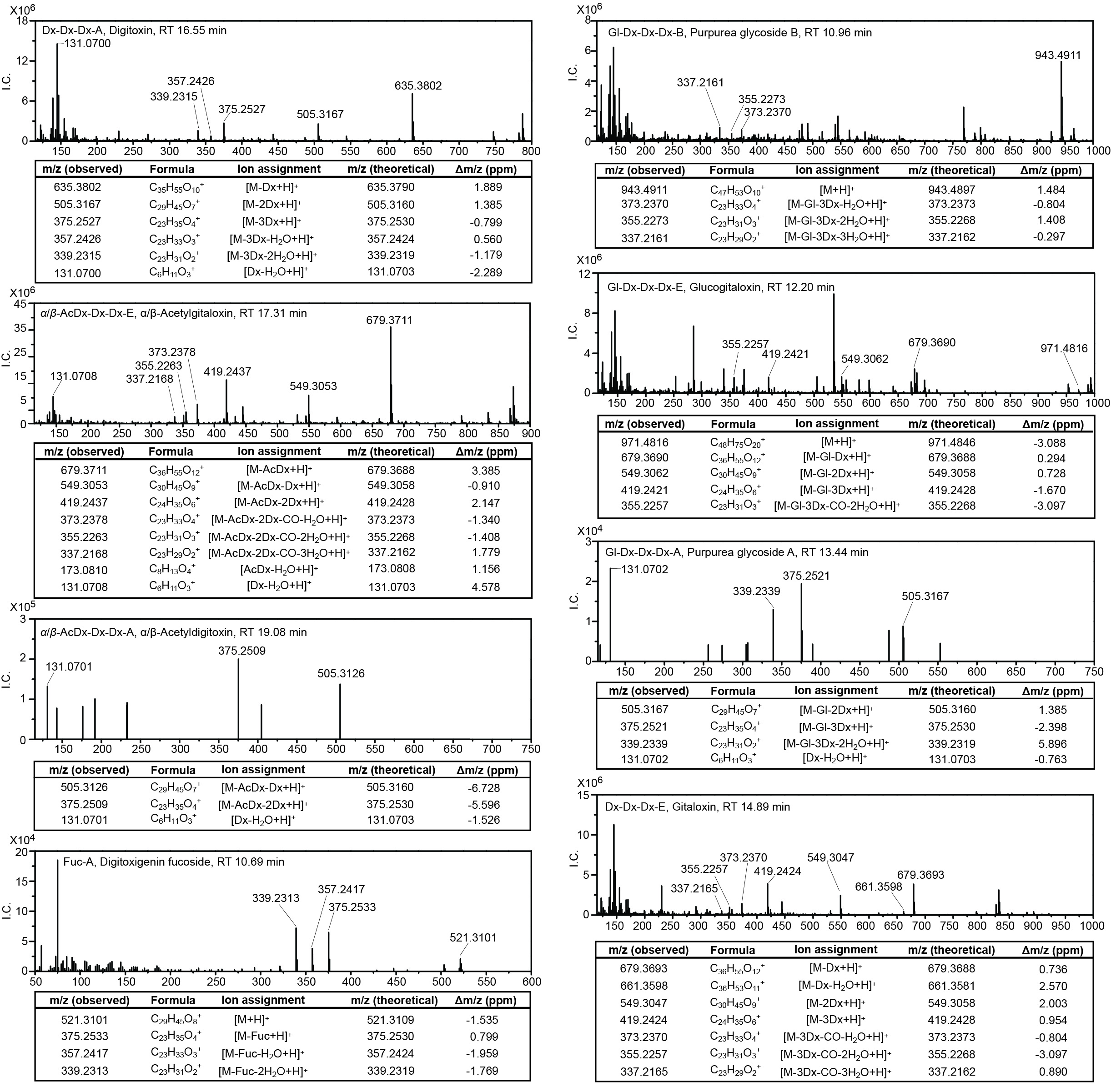

**Fig. S2.** MS^2^ product ion mass spectra from [M+H]^+^ adducts of cardenolides identified in *D.*

*lanata* and *D. purpurea*. I.C.: ion counts; ppm: parts per million; RT: retention time; Aglycones: A, digitoxigenin; B, gitoxigenin; C, digoxigenin; D, diginatigenin; E, gitaloxigenin. Sugars: AcDx, acetyldigitoxose; Dx, digitoxose; Gl, glucose; Dtl, digitalose; Fuc, fucose.
